# Isocall enables scalable transcript identification from long-read RNA-sequencing data

**DOI:** 10.64898/2026.09.08.749180

**Authors:** Egor Dolzhenko, Megan D. Schertzer, Ryan Gossart, Tom Mokveld, Jonathan R. Belyeu, Ales Varabyou, Xinchang Zheng, Elizabeth Tseng, Zev Kronenberg, Mark Chaisson, Gloria Sheynkman, Fritz J. Sedlazeck, Yerbol Z. Kurmangaliyev, Jocelyne Bruand

**Author notes:** These authors contributed equally (Egor Dolzhenko, Megan D. Schertzer, Ryan Gossart, Tom Mokveld, Jonathan R. Belyeu, Ales Varabyou). These authors jointly supervised this work (Fritz J. Sedlazeck, Yerbol Z. Kurmangaliyev, Jocelyne Bruand).

## Abstract

Long-read RNA sequencing directly resolves the full structures of RNA transcripts. Advances in throughput now enable the generation of deeply sequenced cohorts of hundreds of samples, making joint transcript discovery across large datasets possible. However, existing transcript identification methods were designed for small datasets, which limits their applicability at this scale. Here, we present Isocall, a scalable and deterministic computational method for jointly calling transcripts from multiple PacBio long-read RNA sequencing samples. Isocall converts aligned full-length non-concatemer reads into compact per-sample transcript profiles, merges these profiles across samples, and jointly identifies known and novel transcripts supported by reads in the analyzed dataset. Filtering is tunable as presets provide coarse control and individual parameters, including relative abundance and internal priming thresholds, provide fine control. Isocall demonstrated high precision in our accuracy benchmarks, including the 69 transcript WTC11 SIRV spike-in controls, for which Isocall reported 0-2 false-positive transcripts per sample across the three SIRV mixes at default settings. To demonstrate scalability, we applied Isocall to 206 samples, totalling 3.5 billion raw reads, from the Human Pangenome Reference Consortium. After parallelized pbmm2 alignment and Isocall profile, the call step performed joint calling across the entire dataset in 25 minutes, using 1.3 GB of peak memory and 8 threads. Finally, in Genome in a Bottle samples with matched SNP genotypes, splice-site polymorphisms provide an additional measure of call accuracy: Isocall recovered 337 polymorphic splice sites, including a *de novo* donor site in BTN3A1 that corresponds to a complete isoform switch on the mutant allele.

## Introduction

Long-read sequencing (LRS) captures full-length RNA transcript structures, enabling direct characterization of isoform diversity (Sharon et al. 2013; Monzó et al. 2025). As this technology becomes more widespread, RNA LRS studies are rapidly expanding from shallow, single-sample surveys to deeply sequenced cohorts (Al’Khafaji et al. 2023; Reese et al. 2023). These cohort-scale studies are important for characterization of isoform diversity across genes, individuals, and cell types (Reese et al. 2023; Schertzer et al. 2026), interpreting the effects of genetic variation on transcript structure and expression (Glinos et al. 2022; Clavell-Revelles et al. 2025), and understanding the functional consequences of isoform switching and aberrant isoform expression in disease (Aguzzoli Heberle et al. 2025; Shi et al. 2025). As RNA LRS datasets continue to grow in number, sequencing depth, and complexity, there is an increasing need for computational methods that can efficiently analyze transcript isoforms across large cohorts.

There are two broad strategies for identification of transcripts from long-read data: reference-free and reference-based. Reference-free methods reconstruct transcripts by clustering and error correcting the reads without aligning to a genome, as in RATTLE (de la Rubia et al. 2022). These methods are applicable to species without existing genome assemblies or annotations. Reference-based methods align reads to a genome and then extract transcript models from the alignments, either by assembling them, as in StringTie2 (Kovaka et al. 2019), or by grouping reads that share the same splice junctions so that reported transcripts are directly observed rather than assembled, as in FLAIR (Tang et al. 2020), Bambu (Y. Chen et al. 2023), Mandalorion (Volden et al. 2023), and IsoQuant (Prjibelski et al. 2023). Within each broad category, tools differ in many additional decisions, including splice junction correction, transcript start and end annotation, assignment of truncated reads, and filtering thresholds. As a result, tools often report substantially different sets of transcript structures from the same input data (Pardo-Palacios, Wang, et al. 2024). Synthetic spike-in datasets with known transcript structures, such as SIRVs, provide the ground truth needed to evaluate these differences. Filtering thresholds alone can shift precision and sensitivity considerably (Pardo-Palacios, Arzalluz-Luque, et al. 2024).

In addition to accuracy, scalability is becoming an important factor across the existing methods, and computational performance varies widely. In the most recent comparison, StringTie2 (Kovaka et al. 2019) and ISAtools completed analysis of approximately one billion reads at 1.5 GB and 8 GB peak memory, respectively, while IsoQuant, Bambu, and Mandalorion did not complete under the allotted 300 GB of memory (H. Chen et al. 2026). However, both StringTie2 and ISAtools process samples independently, producing sample-specific transcript models that must subsequently be reconciled, complicating comparisons across individuals, families and cohorts. Joint calling instead evaluates the same candidate transcript structures against evidence from all samples, providing a harmonized transcript catalog while retaining sample-level support and improved identification of rare, lowly abundant isoforms, though at a higher computational cost (Jetzinger et al. 2026). Long-read, cohort-scale studies therefore require methods that combine computational scalability with joint, harmonized transcript calling across samples while allowing users to tune precision and sensitivity for their application.

Here, we present Isocall, a computational method for scalable and deterministic joint calling of RNA transcripts from PacBio LRS data. Isocall summarizes aligned reads from each sample into compact transcript profiles and merges these profiles before jointly calling transcript structures. This produces a common set of transcript definitions across the cohort while preserving the sample-specific read support for each transcript. The final ‘call’ step integrates the merged profiles with the provided reference annotation to identify annotated transcripts present in the dataset, discover novel transcripts, and filter likely artifacts. Notably, annotated transcripts not supported by the data are excluded, ensuring that the resulting set of transcripts accurately reflects the transcriptional landscape of the dataset. Isocall inputs aligned full-length non-concatemer (FLNC) reads and outputs a single GTF file of transcript structures and a corresponding matrix of reads supporting each structure in each sample.

## Results

### Isocall performs joint transcript calling across samples

Isocall is a deterministic computational method capable of jointly calling transcripts across hundreds of PacBio LRS RNA samples. Rather than using IsoSeq’s approach of first clustering and collapsing reads into transcript models and then reconciling transcript calls across samples, Isocall aggregates the reads across all samples and only then performs joint transcript calling using all available data. Isocall analysis is separated into four steps: prep, profile, merge, and call (**Figure 1A**). The prep step performs error checking on the reference GTF and converts it into an easy-to-parse format. It needs to be run only once per GTF. During the profile step (**Figure 1B**), aligned BAM files of FLNC reads from each sample are summarized into read bundles. For multi-exon reads, a read bundle contains reads with the same intron chain. The differences in transcript start and end positions are retained within the bundle rather than used to separate the reads. Mono-exonic reads are grouped into bundles based on overlap of their alignments in the reference genome. This representation preserves the transcript structures supported by the reads, while substantially reducing the amount of space required to store this information.

**Figure 1.**
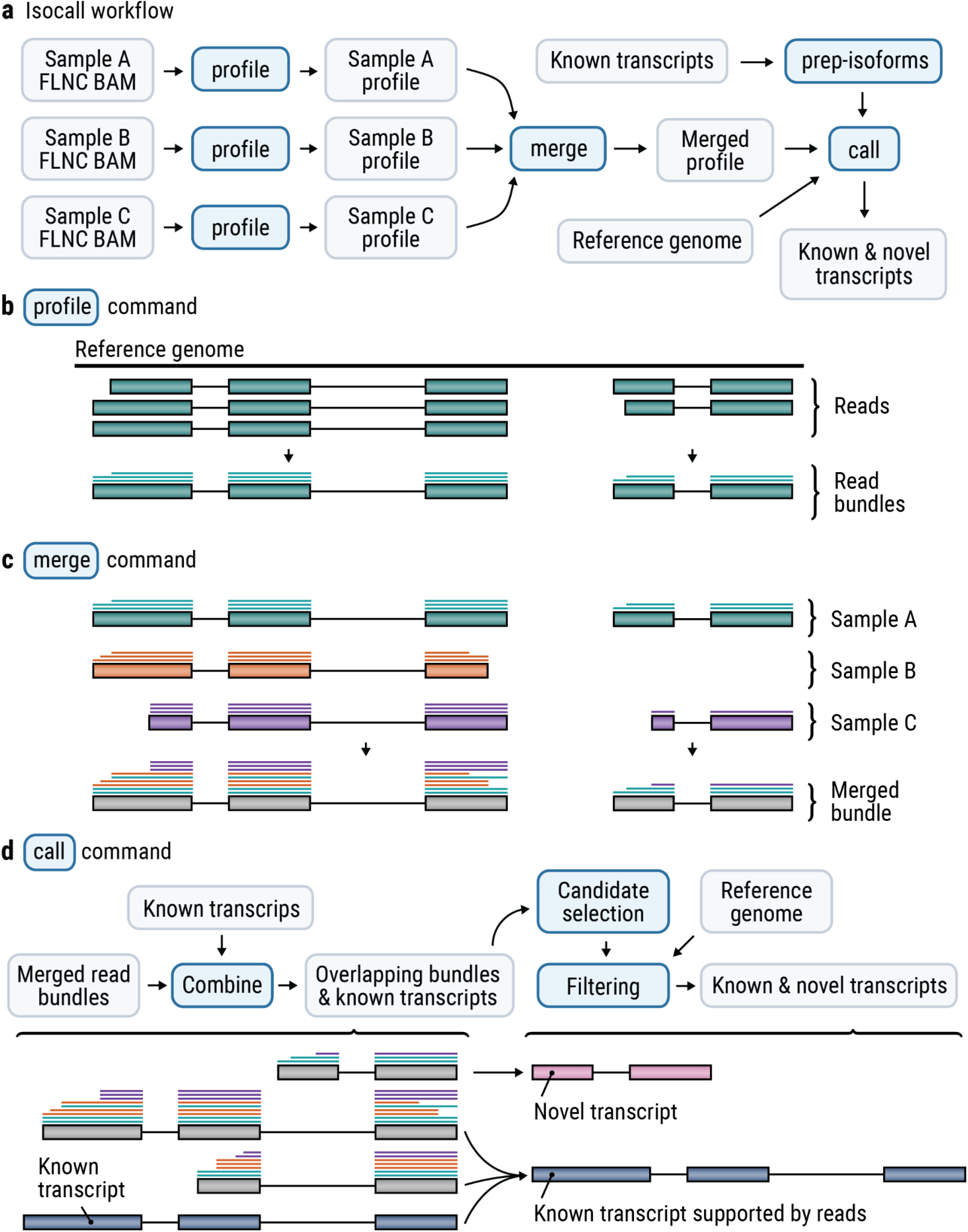
Overview of the Isocall workflow. (A) Full-length non-concatemer (FLNC) reads aligned to the reference genome are converted into per-sample transcript profiles, merged across samples, and jointly analyzed with the known transcript annotations to call known and novel transcripts present in the dataset. (B) Isocall profile extracts exon chains from aligned reads and groups reads with matching intron chains into compact read bundles. (C) Isocall merge combines matching bundles across samples while retaining each read’s sample of origin. Colors depict different samples. (D) Isocall call integrates merged read bundles with known transcripts. Bundles matching an annotated intron chain are reported as known transcripts and the reads they explain are not used for novel transcript discovery; annotated transcripts not supported by reads are not reported. Novel transcripts are then selected from the remaining read bundles by iteratively choosing the candidate with the most compatible reads, and are filtered for internal priming artifacts and low relative abundance.

The merge step (**Figure 1C**) combines compatible read bundles across samples (multi-exonic bundles with the same intron chain or overlapping mono-exonic bundles) while retaining the sample of origin of all reads in a bundle. The call step (**Figure 1D**) then analyzes the merged bundles together with the reference transcript annotations (e.g. GENCODE). Read bundles that match annotated transcript structures provide evidence for presence of known transcripts, while remaining read bundles are used to identify novel transcripts. Isocall applies tunable transcript boundary, internal priming, and relative abundance filters to determine which novel isoforms to report (Methods). These filters can be configured via a single configuration file. For coarser control, Isocall also provides two configuration presets: “default”, which applies the recommended filters and thresholds to produce a high-confidence transcript set, and “yolo”, which disables read bundle downsampling and minimizes filtering to maximize sensitivity.

### Isocall provides tunable precision at low computational cost

Isocall replaces the ‘cluster’ and ‘collapse’ steps of the IsoSeq workflow, calling transcripts from reference alignments rather than from de novo clustering of FLNC reads. To compare the performance of Isocall (default and yolo presets) and IsoSeq, we analyzed WTC11 PacBio LRS datasets containing SIRV E0, E1, and E2 spike-ins, which represent 69 transcripts across seven SIRV genes (Wissel et al. 2026), and scored endogenous calls against TUSCO, a control set of 46 single-isoform genes (Liu et al. 2026). Isocall with default settings showed substantially higher precision than IsoSeq, calling a median of 0, 2, and 0 false positives across SIRV mixes E0, E1, and E2 compared with 99.5, 582, and 231, respectively (**Figure 2A**). However, this higher precision came at the cost of more false negatives, with a median of 1.5, 10, and 14 for Isocall compared with 1, 1, and 3 for IsoSeq. This tradeoff is expected. The Isocall default preset prioritizes precision to produce a high-confidence transcript set, while the yolo preset applies minimal filtering to prioritize sensitivity, recovering nearly all true transcripts (median of 0, 1, and 2 false negatives) at the cost of many more false positives (263, 965, and 547). Within TUSCO, false positive patterns were similar though less pronounced, and false negatives were comparable across all three.

**Figure 2.**
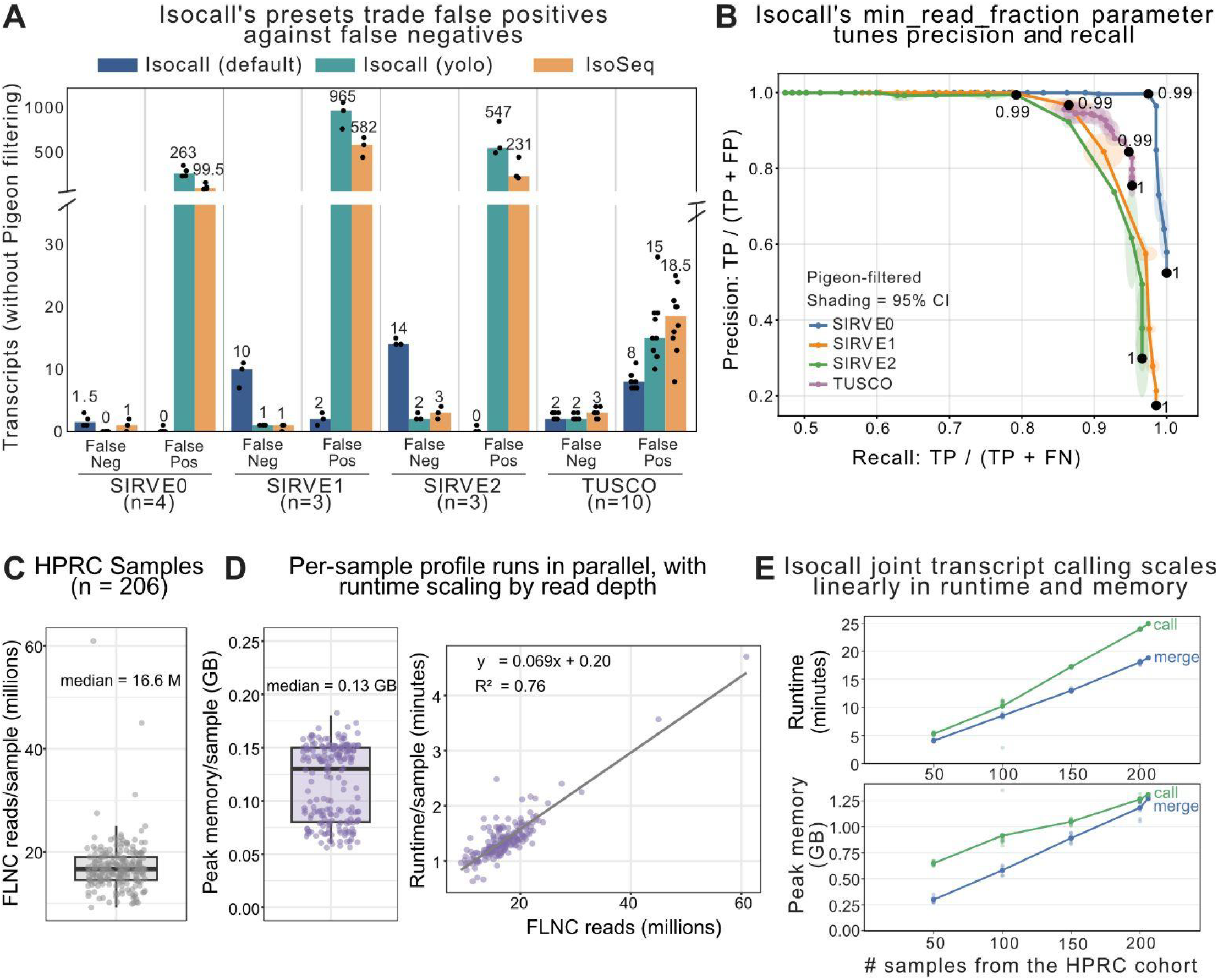
Isocall provides tunable precision at low computational cost. (A) False negatives and false positives, without Pigeon filtering, called by Isocall (default and yolo presets) and IsoSeq across SIRV spike-in mixes E0, E1, and E2 and the TUSCO endogenous gene set. Each of the 10 WTC11 samples contains one SIRV mix, and all 10 were scored against TUSCO. Bars show the median across samples, with the median value printed above each bar. Points show the individual samples. (B) Pigeon-filtered precision and recall across `min_read_fraction` values (0.85 to 1.00) for SIRV mixes and the TUSCO gene set. Shading indicates the 95% confidence interval across samples. Black points mark the values used by default (0.99) and yolo (1) presets. Because (A) is unfiltered and (B) is filtered, false positive counts in (A) are not directly comparable to the precision values in (B). (C) Distribution of FLNC reads per sample, in millions, across the 206 HPRC PacBio RNA LRS samples. Box shows the median and interquartile range and points show individual samples. (D) Peak memory and runtime for the Isocall profile step, which runs independently on each HPRC sample. Left: peak memory per sample, with the box showing the median interquartile range and points showing individual samples. Right: runtime per sample versus FLNC read depth. Runtime scaled at approximately 4 seconds per million reads. (E) Runtime and peak memory for the Isocall merge and call steps, which run jointly across all samples. Random subsets of 50, 100, 150, and 200 samples were drawn ten times each without replacement, together with the full cohort of 206 samples, which was measured once. Points show individual draws.

The Isocall presets demonstrate the two extremes of the precision-sensitivity tradeoff, while individual parameters allow finer control. For example, `min_read_fraction` sets the fraction of each gene’s reads to retain, discarding the lowest-support read bundles. It can be tuned from 0.85 for higher precision toward 1 for maximum recall (**Figure 2B**). This parameter alone accounts for most of the differences between the default (0.99) and yolo (1.00) presets. At 0.99, precision ranged from 0.84 to 0.996 and recall from 0.79 to 0.98. At 1, where all read bundles are retained, precision dropped to 0.17-0.76 while recall increased to 0.95-1. The appropriate value depends on the user application, lowering the parameter increases confidence while raising it toward 1 will recover more transcripts.

To assess computational cost at cohort-scale, we applied Isocall with default settings to 206 PacBio RNA LRS samples from the Human Pangenome Reference Consortium (HPRC), with a total of 3.5 billion raw FLNC reads (median = 16.6 million, **Figure 2C**) (Liao et al. 2023). The profile step runs independently on each sample, so its cost is unaffected by cohort size. However, runtime did scale with read depth at approximately 4 seconds per million reads, with the deepest sample (60 million reads) completing in 4.7 minutes (**Figure 2D**). Peak memory was independent of read depth, with a median of 0.13 GB per sample. The merge and call steps run jointly across all samples and therefore scale with cohort size. Across subsets of 50-206 samples, runtime increased approximately linearly, from 4.1 to 18.8 minutes for merge and 5.3 to 24.9 minutes for call (**Figure 2E**). Extrapolating from the observed linear trend, we expect merge and call to scale to roughly 90 and 120 minutes, respectively, on a cohort of 1,000 samples of similar read depth. Peak memory did not exceed 1.4 GB throughout both steps when running on the full cohort of 206 samples. Together, these results show that Isocall enables tunable precision and sensitivity in transcript discovery while substantially reducing runtime and memory requirements.

### Isocall discovers changes in splicing patterns driven by genetic variation

The SIRV benchmark establishes that Isocall precisely identifies ground truth transcript structures, but these are synthetic controls that are already annotated. As a complementary source of ground truth, we leverage genetic variation, since variants that disrupt or create canonical GT-AG splice site dinucleotides alter transcript structure in predictable ways (Burset et al. 2000; Kurmangaliyev et al. 2015, 2013).

Genome in a Bottle (GIAB) provides LRS with matched genotypes from three individuals within two family trios: the Ashkenazi mother (HG004) and son (HG002) and the Chinese son (HG005; **Figure 3A**) (Zook et al. 2016). Joint transcript calling with Isocall across these three GIAB samples detected 48,464 reference transcripts and 127,438 novel transcripts (**Figure 3A**). At the level of individual splice sites, we detected 312,431 reference sites and 58,762 novel sites, with a near-even split between donors and acceptors (**Figure 3B**). We then identified splice sites overlapping single-nucleotide polymorphisms (SNPs) in canonical GT-AG dinucleotides, filtering to sites where at least two individuals carry different combinations of functional and non-functional alleles. For example, a homozygous functional acceptor (AG/AG) in one individual and a heterozygous site (AG/TG) in another. In total, we identified 118 reference and 219 novel polymorphic splice sites (**Figure 3B**).

**Figure 3.**
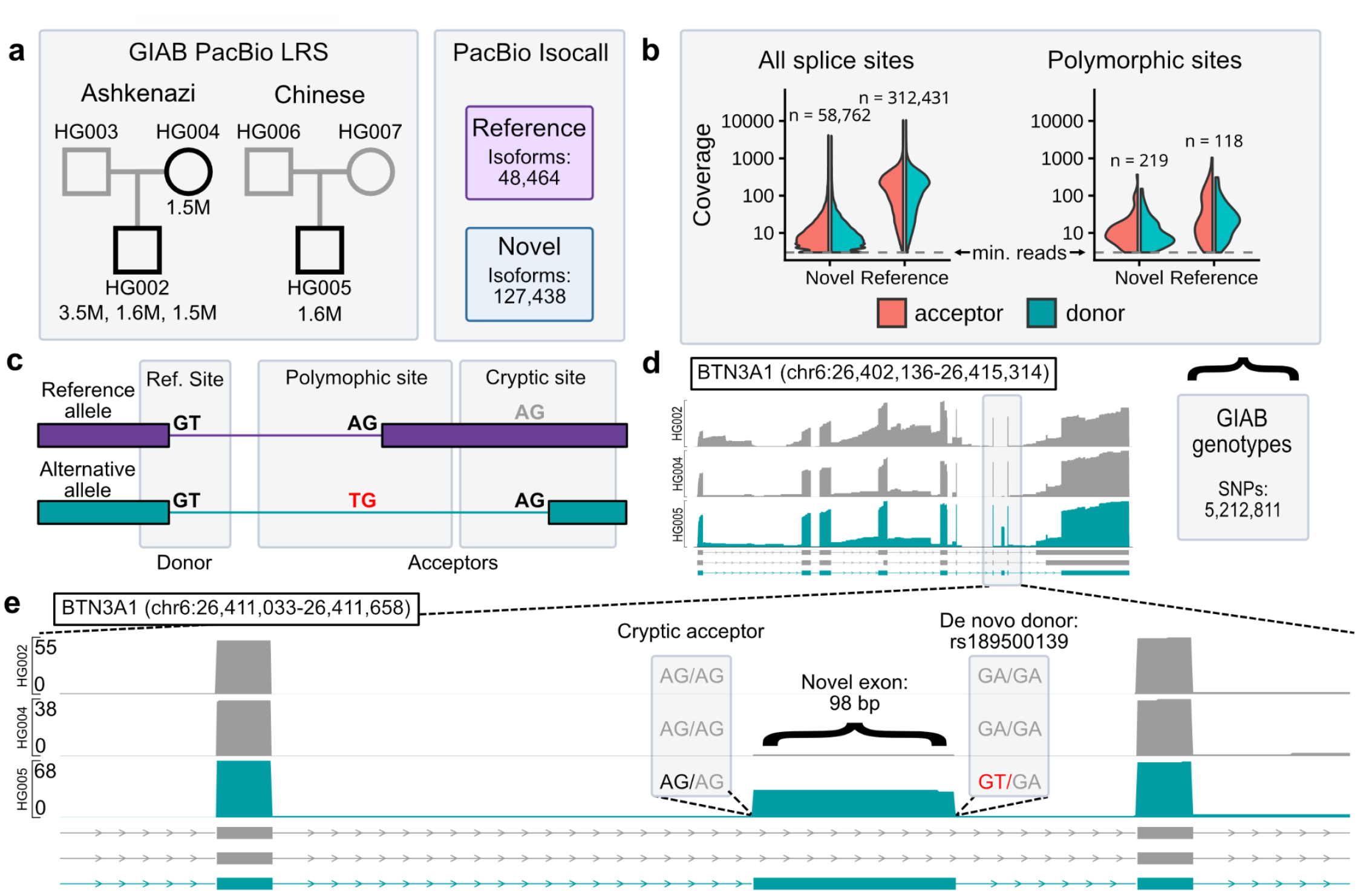
Isocall detects outcomes of splicing mutations. (A) Summary of the GIAB long-read dataset and Isocall transcript calls. The dataset includes five libraries for three individuals (3x HG002, HG004, HG005). Library sizes are indicated (1.5 to 3.5 million reads per sample). (B) Cumulative coverage of splice sites across the whole dataset. (C) Schematic of a polymorphic splice site in which a single-nucleotide polymorphism (SNP) disrupts an acceptor site and causes activation of a downstream cryptic site that is not used by the reference allele. In total, we detected 337 polymorphic splice sites. (D-E) De novo splice site in BTN3A1. Plots show the coverage for each individual and representative transcripts for the whole locus (D) and the affected region (E). The heterozygous A-to-T variant in HG005 creates a de novo donor site and activates an upstream cryptic site that is present in all individuals. As a result, a de novo 98 bp long exon is inserted into the coding region of the gene, potentially disrupting the encoded protein. Half of the reads in HG005 include this exon, consistent with a complete switch of isoforms expressed by the mutated allele.

Most novel splice sites detected by Isocall had low coverage, suggesting that most novel transcripts are also lowly expressed in these samples (**Figure 3B**). Furthermore, both reference and novel polymorphic splice sites had low coverage. This is consistent with previous observations that polymorphic splice sites detected in healthy individuals are primarily used in lowly expressed transcripts and have minor effects on protein structure and gene function (Kurmangaliyev et al. 2013).

Some polymorphic sites were used at high frequencies and likely represent a switch in major isoform between alleles. One such example was detected in *BTN3A1*, a gene involved in T cell activation and proliferation (Messal et al. 2011) (**Figure 3D**). The reference genome and both Ashkenazi samples, HG002 and HG004, carry a non-functional allele at this donor site (GA/GA), while HG005 is heterozygous for a functional splice site (GT/GA). The functional allele frequency of this SNP (<1%) (1000 Genomes Project Consortium et al. 2015) and whole-genome alignments with other primates (Casper et al. 2026) suggest that the non-functional variant is ancestral and that this case represents the de novo creation of a new splice site.

Isocall detects a transcript for *BTN3A1* which includes a novel exon using this de novo splice site. The acceptor site of this exon is a cryptic splice site that is present in all three individuals but is only activated in HG005 (**Figure 3E**). About half of the reads in this sample include this exon (33 of 68), indicating that the novel exon is constitutively used by the functional allele and represents a complete switch of the major isoform by this mutation. The exon is 98 bp long and therefore predicted to cause a frameshift in the coding sequence, altering the downstream protein sequence. This example demonstrates that Isocall can detect expected changes in splicing patterns driven by genetic variation.

## Discussion

Isocall is designed to enable scalable transcriptomics studies involving hundreds of samples. While most existing transcript identification methods are designed to analyze one or few samples at a time, Isocall calls transcripts by analyzing all filter-passing reads across all samples at the same time, thus providing a harmonized transcript catalog across multiple samples.

One of the clearest improvements of Isocall over the IsoSeq workflow is specificity. On SIRV spike-ins, where every called transcript that is not one of the 69 known SIRV transcripts is a false positive, Isocall at default settings reported a median of at most two false positives per sample across the three mixes, compared with hundreds for the IsoSeq cluster / collapse workflow. The gap between the methods widens on more complex samples: the E1 mix, with a four-fold concentration range, produced 582 false positives per sample with IsoSeq and two with Isocall.

Filtering steps play a crucial role in Isocall performance, as can be seen in the large difference in false positive rates between the default and the yolo mode. This is especially evident in structurally complex transcripts in SIRV benchmarks. In contrast, for the simpler single-isoform control genes included in the TUSCO benchmark false negatives were comparable across Isocall configurations (because the filters had less to remove). The high sensitivity analysis of the yolo mode could thus be useful in discovery studies that may use different, study-specific filtering criteria. Isocall’s efficiency is also worth noting. Joint transcript calling across the HPRC cohort of 206 samples totalling 3.5 billion reads completed in just 25 minutes using 1.3 GB of peak memory while utilizing just 8 threads.

An important advantage of joint transcript calling is that transcript evidence is evaluated consistently across the entire cohort rather than independently within each sample. This is particularly relevant for low-abundance isoforms, for which individual samples may contain insufficient read support to pass transcript-calling thresholds. When the same transcript structure is observed across multiple samples, joint calling can preserve this distributed evidence and incorporate the transcript into a common cohort-level catalog while retaining sample-specific read support. By contrast, independent calling can produce sample-specific transcript sets that differ because of local read depth and filtering thresholds and must subsequently be reconciled. Joint calling therefore provides a natural framework for harmonizing transcript definitions across individuals, families and cohorts, which is increasingly important as long-read RNA-sequencing studies expand to population scale. Importantly, annotated transcripts for which no supporting reads were observed are not reported by Isocall. This is an improvement over approaches that carry the full set of reference annotations through to quantification, because transcripts absent from the dataset will still absorb ambiguously assigned reads.

Our results show that joint read-level analysis of transcriptomics datasets can be performed efficiently even for datasets containing hundreds of samples. Isocall’s high computational efficiency and determinism allow creating reproducible cohort-scale resources.

## Methods

### Isocall Tool

Isocall jointly calls transcripts across multiple samples from reference-aligned FLNC reads. The workflow comprises four stages: 1) preparation of a database of known transcripts from reference annotation, 2) generation of per-sample transcript profiles, 3) merging of profiles across samples, and 4) joint calling, annotation, filtering, and quantification of known and novel transcripts. Unless otherwise specified, default calling parameters were used.

### Isocall prep-isoforms: Database of known transcripts

A reference annotation (e.g., GENCODE) in GTF format is converted into a compact database of transcript models. Exon coordinates per transcript are condensed into a single line representation, sorted by genomic position, and associated with their corresponding gene identifiers and gene symbols. The resulting database format enables efficient exon chain matching between known transcripts and long reads in the downstream “isocall call” command.

### Isocall profile: Per-sample transcript profiles

For each sample, Isocall profile inputs an aligned BAM file and retains only primary alignments where at least 95% of the read sequence is aligned to the reference genome. Each retained alignment is converted to an exon chain by treating reference-skip (N) CIGAR operations as introns, while ignoring any mismatches, insertions, and deletions within aligned exons.

Reads are then grouped into read bundles. Multi-exonic reads with identical splice junction chains are collapsed into a single bundle, ignoring TSS and TES coordinates so that reads differing only in their start and end positions are still bundled. Similarly, mono-exonic reads are collapsed when their genomic intervals overlap on the same strand. The output of this command is a gzip-compressed single-sample profile containing sample metadata, chromosome lengths, and read bundles. Each bundle is stored as a single row, and bundle start and end are defined by the leftmost read start and rightmost read end. To store bundles compactly, bundle exon coordinates are encoded as offsets from the bundle start rather than as absolute genomic coordinates.

### Isocall merge: Joint transcript profiles

Single-sample transcript profiles from Isocall profile are merged together into a joint transcript profile. The read bundles are merged across samples using the same splice junction chain matching (multi-exon) and genomic interval overlap (mono-exon) rules used during per-sample profiling. Merging is performed in genomic coordinates, with exon offsets recomputed against the merged bundle start. The final column of the merged output retains comma-separated read information formatted as `<start>_<end>_<readcount>_<sample-index>`, where the read start is offset from bundle start and read end is offset from the start of the last exon.

### Isocall call: Joint transcript calling

Isocall next groups read bundles by gene locus. To do this, it builds a graph whose nodes are the exons observed in the merged read bundles together with the exons of annotated transcripts. Two exons are connected when they overlap, when they are splice-adjacent within a read bundle or annotated transcript, or when they belong to the same annotated gene. By default, read bundles with total support below 3 reads are excluded from the analysis (`min_reads_per_isoform` parameter). To control the processing speed of highly expressed loci, bundles assigned to each gene are ranked by total read support and then only the top bundles containing at least 99% of reads (`min_read_fraction` parameter) are retained, subject to a maximum of 10,000 best-supported read bundles per gene (`max_bundles_per_gene` parameter). During this downsampling step, read bundles overlapping multiple genes are assigned to the highest-expressed overlapping gene. Known multi-exonic transcripts are identified by identical splice junction chain matching to the reference database (from `prep-isoforms` command). Known multi-exonic transcripts are detected by exact strand-specific intron chain matching, irrespective of terminal exon boundaries. A known mono-exonic transcript is detected when a retained mono-exon read bundle is fully contained within its annotated interval. Because bundle filtering precedes identification, the contributing bundle must have total cohort support of at least three reads under the default settings. After known transcripts have been identified, reads explained by those transcripts are removed. For bundles that match known transcripts, the start and end are defined by their annotated coordinates unless optional known-end extension is enabled (`extend_known_ends` parameter).

To call novel multi-exonic transcripts, candidate splice-junction chains are taken from remaining read bundles. For each multi-exonic candidate, internal splice boundaries are taken from the bundle. The 5′ and 3′ boundaries are set after excluding the outermost 10% of observations at each end (`terminal_trim_fraction` parameter). Candidates are then selected using a greedy support-maximization procedure that iteratively chooses the candidate supported by the largest number of compatible reads and removes reads explained by that candidate before selecting the next candidates.

Remaining mono-exon reads are processed separately using strand-aware clustering of their 3′ ends. Reads are sorted by 3′-end position, and adjacent observations separated by at most 20 bp are placed in the same cluster. A cluster is called when it has support from at least 10 reads. Its 3′ boundary is the modal read-end position, and its 5′ boundary is determined after trimming the outermost 5% of weighted 5′-end observations. Overlapping candidate intervals are subsequently merged. On chrM, novel discovery is skipped and known transcripts are passed through. Novel transcripts assigned to an annotated gene are named “`<gene_id>|T<N>`“. Intergenic loci receive coordinate-derived gene identifiers, and their transcripts are named by appending “`|T<N>`“. Novel transcripts that overlap multiple genes are labeled as putative fusions when there are adjacent exons that are assigned to different annotated genes; remaining ambiguous cases are assigned to the gene with the highest exon-chain Jaccard overlap.

### Additional filtering of novel transcripts

Two post-calling filters are applied to novel transcripts. First, an internal priming filter examines the 20-bp sequence downstream of the transcript 3’ end (`window` parameter). A novel transcript is flagged if this sequence has A-content >0.60 or contains at least 6 consecutive As (`fraction` and `run_length` parameters). Flagged transcripts are rescued if their transcription termination site lies within 25 bp of an annotated transcript end site from the same gene (`rescue_distance` parameter). Second, a relative abundance filter suppresses low-abundance novel transcripts that are structurally similar to a more abundant transcript. Structural distance is computed from internal splice-boundary differences and non-overlapping exonic segments (terminal boundary differences are not penalized). The filter is applied only for distances of at most 20 (`max_distance` parameter), using a piecewise threshold of 0.1^floor((d-1)/5) (`abundance_base` parameter).

After filtering, Isocall outputs a table reporting the number of supporting reads for each transcript in each sample. A multi-exonic read is assigned to every reported transcript with the same strand-specific splice junction chain, whereas a mono-exon read is assigned to every reported single-exon transcript that fully contains it. Consequently, reads compatible with multiple transcripts are counted for each compatible transcript rather than fractionally allocated.

### Increased sensitivity preset

Isocall call supports running with a preset parameter configuration called `yolo` mode, intended for increased sensitivity relative to default configuration. This configuration sets the `min_read_fraction` to 1.0 to disable read bundle downsampling, disables filtering of putative internal priming artifacts via poly-A identification, and disables the relative abundance filter. These options increase the sensitivity of Isocall call commands and decrease specificity.

### Benchmarking and evaluation

#### Benchmarking PacBio Isocall versus IsoSeq on SIRV and TUSCO datasets

Isocall precision and recall were characterized against the Lexogen SIRV spike-in gene set and well-characterized TUSCO gene set in PacBio LRS of undifferentiated WTC11 human-induced pluripotent stem cells (Wissel et al. 2026). Ten datasets were used for testing: four with the Lexogen SIRV E0 mix, three with SIRV E1 Mix, and three with the SIRV E2 Mix. The SIRV mixes represent 69 total transcripts across seven SIRV genes, with concentration ratios of equal representation in the E0 Mix (four datasets), ranging from 1/2x - 2x in E1, and ranging from 1/32x - 4x in E2. The TUSCO curated gene set (Liu et al. 2026) includes 46 universally expressed single-isoform genes found in all human tissue types.

Reads were aligned to a combined reference of human GRCh38 and SIRV with pbmm2 v26.2.99 (using the ‘ISOSEQ’ preset). Datasets were standardized by down-sampling SIRV-aligned reads to 1 million per dataset, while non-SIRV reads were all included. Isoforms were characterized twice with Isocall v1.3.0-c082284 using the two presets, default and ‘yolo’. Transcript discovery was also performed from the same data with IsoSeq (IsoSeq data retrieved from https://downloads.pacbcloud.com/public/dataset/Kinnex-bulk-2023-WTC11/). Results from Isocall and IsoSeq were each classified with Pigeon v26.2.99, but not filtered.

Transcripts from each group (Isocall default, Isocall ‘yolo’, and IsoSeq3) were considered true positives if classified by Pigeon as a full-splice match to a known isoform and false positives if classified as novel. Transcripts in the reference annotation that were not called were considered false negatives. **See Figure 2A**. Incomplete splice match transcript calls were not considered in precision and recall metrics.

To assess the contribution of the `min_read_fraction` parameter, Isocall was additionally run across values from 0.85 to 1.00 with all other settings kept at their defaults. These results were filtered and classified with Pigeon v26.2.99 with poly-A motif scoring enabled and minimum reference transcript length set to 100 bp. Precision and recall were calculated as discussed above. **See Figure 2B**.

#### Measuring performance on the HPRC dataset

To measure Isocall’s performance on a representatively large cohort, we used 206 publicly available deep-sequenced Human Pangenome Reference Consortium (HPRC) PacBio Kinnex RNA samples (Liao et al. 2023) (HPRC Data Release 2, https://humanpangenome.org/hprc-data-release-2/) and aligned to GRCh38 with pbmm2. Across the cohort, each sample carried a mean of 17.2 M aligned reads (SD 5.0 M; median 16.8 M). Transcript calling was guided by the GENCODE v49 annotation, from which the known-transcript model was built once with isocall prep-isoforms. Over the full cohort, Isocall called 1,788,390 transcripts, to which 76.3% (SD 7.7%) of each sample’s aligned reads were assigned.

Isocall v1.3.0 with default parameters were used for all performance evaluation. Isocall’s four stages scale differently with cohort size: prep-isoforms depends only on the annotation; profile runs independently per sample; and merge and call are cohort-wide. To characterise the growth of the cohort-dependent stages, we drew random subsets of 50, 100, 150 and 200 samples, together with the full cohort of 206 samples. Each size was drawn ten times without replacement, except the full cohort, for which only one subset exists and which was therefore measured once, giving 41 subsets and 164 stage measurements. Isocall prep-isoforms was measured once and reused. Isocall profile, being independent of cohort composition, was measured once per sample, and its cost for a draw obtained by summing the per-sample timings over the drawn samples. Isocall merge and call were re-executed for every draw.

Elapsed wall-clock time and memory usage were recorded for each stage executed with 8 threads and a 32 GB memory request. All measurements were made on a single compute node (two 64-core x86_64 sockets, 128 CPUs at one thread per core, 515 GB RAM, local NVMe) held under an exclusive Slurm allocation, with the workflow restricted to one active allocation at a time to prevent competing resource usage.

#### Analysis of the GIAB dataset

Long-read alignments and genotypes were downloaded from the Genome in a Bottle (GIAB) project repository (Zook et al. 2016). A dataset-specific transcriptome was called using Isocall v1.3.0 with the default configuration, GRCh38 as the reference genome, and GENCODE v39 as the reference gene annotation (Mudge et al. 2025). A custom python script was used to parse per-sample transcript profiles and pull junction counts.

We merged the VCF files for HG002, HG004, HG005 and extracted SNPs that overlapped with canonical splicing nucleotides at the first two or last two bases of each intron in the dataset-specific annotation generated by Isocall. Next, we identified polymorphic splice sites where the splice site genotypes differed between at least two individuals. VCF filtering and coverage plot generation used the following Bioconductor packages (release 3.22): GenomicRanges, GenomicFeatures, VariantFeatures, rtracklayer, and trackviewer (Huber et al. 2015).

## Declarations

### Data availability

All datasets analyzed in this study are publicly available. SIRV spike-in data were obtained from the public PacBio Kinnex-bulk-2023-WTC11 dataset (https://downloads.pacbcloud.com/public/dataset/Kinnex-bulk-2023-WTC11/). The TUSCO reference gene set is distributed with SQANTI3 (https://github.com/ConesaLab/SQANTI3). HPRC PacBio Kinnex RNA data are available under NCBI BioProject PRJNA701308 (SRA study SRP305758) and through HPRC Data Release 2 (https://humanpangenome.org/hprc-data-release-2/). GIAB long-read alignments and genotypes were obtained from the GIAB project repository (https://www.nist.gov/programs-projects/genome-bottle).

### Method availability

Isocall is public and distributed under PacBio’s license on GitHub: https://github.com/PacificBiosciences/isocall

